# Rapid self-selecting and clone-free integration of transgenes into engineered CRISPR safe harbor locations in *Caenorhabditis elegans*

**DOI:** 10.1101/2020.05.26.117390

**Authors:** Zachary C. Stevenson, Megan J. Moerdyk-Schauwecker, Brennen Jamison, Patrick C. Phillips

## Abstract

Precision genome editing for model organisms has revolutionized functional analysis and validation of a wide variety of molecular systems. To date, the capacity to insert transgenes into the model nematode *Caenorhabditis elegans* has focused on utilizing either transposable elements or CRISPR-based safe harbor strategies. These methods require laborious screening processes that often result in false positives from heritable extrachromosomal arrays or rely on co-CRISPR markers to identify likely edited individuals. As a result, verification of transgene insertion requires anti-array selection screening methods or extensive PCR genotyping respectively. These approaches also rely on cloning plasmids for the addition of transgenes. Here, we present a novel safe harbor CRISPR-based integration strategy that utilizes engineered insertion locations containing a synthetic guide RNA target and a split-selection system to eliminate false positives from array formation, thereby providing integration-specific selection. This approach allows the experimenter to confirm an integration event has taken place without molecular validation or anti-array screening methods, and is capable of producing integrated transgenic lines in as little as five days post-injection. To further increase the speed of generating transgenic lines, we also utilized the *C. elegans* native homology-based formation of extra-chromosomal arrays to assemble transgenes *in-situ*, removing the cloning step. We show that complete transgenes can be made and inserted into our split-selection safe harbor locations starting from PCR products, providing a clone-free and molecular-validation-free strategy for single-copy transgene integration. Overall, this combination of approaches provides an economical and rapid system for generating highly reproducible complex transgenics in *C. elegans*.

## Introduction

The introduction of transgenes is a staple in the molecular biologist’s toolkit, with a broad range of utilities including expression of individual variants, ectopic expression of tagged native genes, and the addition of genes from other species. Injection of double-strand DNA into the *Caenorhabditis elegans* gonad arm generally results in assembly of these fragments via regions of microhomology, leading to the formation of extrachromosomal arrays (Stinchcomb *et al.* 1985; Mello *et al.* 1991). These extrachromosomal array structures can be in excess of 1 Mbp and contain up to hundreds of copies of the injected gene (Mello *et al.* 1991; Woglar *et al.* 2019). Extrachromosomal arrays are not stably inherited—either between cells within an individual or between generations—and have variable expression levels, which can be problematic depending on the biological question. To avoid the stochastic element of array expression, it is often desirable to integrate transgenes. Historically in *C. elegans,* microparticle bombardment (Praitis *et al.* 2001) and ultraviolet light exposure (Evans 2006) or gamma exposure (Mello and Fire 1995). have been used to integrate transgenes randomly. However, these methods are less than ideal as: integration usually results in multiple copies, which can impact expression; the integration location is random which can disrupt native genes expression or insert the transgene integrate into regions prone to transcriptional silencing; and both methods require expensive and specialized equipment to create transgene integrations and to identify the loci of insertion. More recent approaches have utilized transposon-based integration methods such as MosSCI and miniMos (Frøkjær-Jensen *et al.* 2008, 2014), which use a transposon to create a double-strand break, allowing for single transgene integration into a predefined or random region respectively. Recently, CRISPR/Cas9 techniques have been adopted for transgene integration. This includes generalized methods for transgenic cargo insertion, either with or without a selective marker such as Hygromycin B resistance (Radman *et al.* 2013) or a self-excising cassette (SEC) (Dickinson *et al.* 2015; Kasimatis *et al.* 2018). More specialized CRISPR strategies, such as the SKI LODGE method, facilitates tissue-specific expression by splitting the coding and promoter element (Silva-García *et al.* 2019). A particular advantage of this latter strategy is that it introduces modular transgene integration, allowing for a more straightforward integration into a backbone of standard genetic elements that are pre-integrated within a safe harbor location.

Regardless of whether a transposon or CRISPR/Cas9 strategy is used, integration of the transgene by homology-directed repair (HDR) is generally inefficient compared to non-homologous end joining (NHEJ) (Frøkjær-Jensen *et al.* 2008; Dickinson *et al.* 2013, 2015; Ward 2015). Both MosSCI and CRISPR show approximately the same integration efficiencies which varies greatly depending on the transgene. CRISPR and MosSCI also require robust screening methods to identify the rare correct transgene integration. Co-CRISPR simultaneously targets a second gene to generate a visible dominant phenotype (e.g., *dpy-10)* (Arribere *et al.* 2014), thereby allowing identification of a sub-population with active Cas9 expression and genome targeting. This enriched population must then be further screened, generally by PCR and Sanger sequencing, to identify correct integration events at the desired locus. In the case of selectable genes, the transgene generates a visible phenotype displayed not only by integrants but also individuals with heritable extrachromosomal arrays. This is because the injected donor homology contains a fully functional copy of the selectable gene. Distinguishing extrachromosomal arrays from integration events requires anti-array selection techniques such as heat shock induction of peel, a toxic transgene (Seidel *et al.* 2011; Frøkjær-Jensen *et al.* 2012) and visual screening for loss of an additional gene used to mark the array (e.g., fluorescent protein or *rol-6).* These methods are imperfect and molecular methodologies such as genotyping PCRs must be used to verify genuine integrations. Currently, no method is available for *C. elegans* that provides integration-specific selection of transgenes. Such an approach would reduce the labor required to generate transgenics by eliminating false positives caused by extrachromosomal array formation, fitting into the category of a “screen from heaven” where only the desired transgenic integrant is alive on the petri dish (Jorgensen and Mango 2002).

In most cases, transgenes must be cloned into plasmids with homology arms matching the targeted genomic region for single-copy integration. This process requires laborious cloning strategies for each desired transgene. *C. elegans* can recombine fragments with microhomology and express resulting transgenes in an array (Mello *et al.* 1991; Kemp *et al.* 2007). Others have tested this strategy to create a donor homology for transgene integration. For example, Paix *et al.* (2016) and Philip *et al*. (2019) attempted to overcome the cloning obstacle by integrating transgenes with overlapping PCR fragments that, once recombined *in-situ,* should produce a functional gene. However, neither method provides direct selection for the transgene integration. As such, depending on the design, array formation can provide false-positives, increasing the difficulty of identifying a correct assembly and integration—a notable complication for current ‘clone-free’ strategies.

Here, we present a novel transgene integration strategy that utilizes a custom-designed safe harbor location to eliminate many of the steps required to go from concept-to-integrated transgene. Our approach removes the selective advantage from the array and selects only for the integration event by splitting the coding sequence for Hygromycin B resistance. Additionally, we show the cloning stage can be bypassed in this system, utilizing the worms’ native homology mediated repair to clone our transgene *in-situ*. Coupling these methods can reduce the labor and time required to produce a transgenic nematode, allowing the experimenter to go from PCR-to-integrated transgene in approximately one week.

## Materials and Methods

### Strains and growth conditions

Bristol N2-PD1073 (Yoshimura *et al.* 2019) and the derived strains PX692, PX693, PX694, PX695, PX696, and PX697 (Table S1) were maintained on NGM-agar plates seeded with OP50 or HB101 *Escherichia coli* at 15°C unless otherwise noted.

### Molecular biology

All plasmids unique to this publication are listed in Table S2, and all primers used in this study are listed in Table S3. All-in-one-plasmids encoding both Cas9 and the desired sgRNA were created by site-directed mutagenesis of pDD162 (Addgene #47549) (Dickinson *et al.* 2013) using the Q5 site-directed mutagenesis kit (NEB) per manufacturer directions. Guide and Cas9 sequences were confirmed by Sanger sequencing. The guide targeting Chromosome II:8420188-8420210 has been previously described, and the constructed plasmid (pMS8) is equivalent to pDD122 (Addgene #47550) (Dickinson *et al.* 2013). Synthetic guide sites utilized in the landing pads were based on guides previously shown to be highly efficient in other species or generated based on predicted efficiency scores (Table 1). Predicted off-target effects were determined using the method of (Doench *et al.* 2016) while predicted on-target efficiency was calculated using Sequence Scan for CRISPR (Xu *et al.* 2015) and the method of (Hsu *et al.* 2013).

**Table 1:**
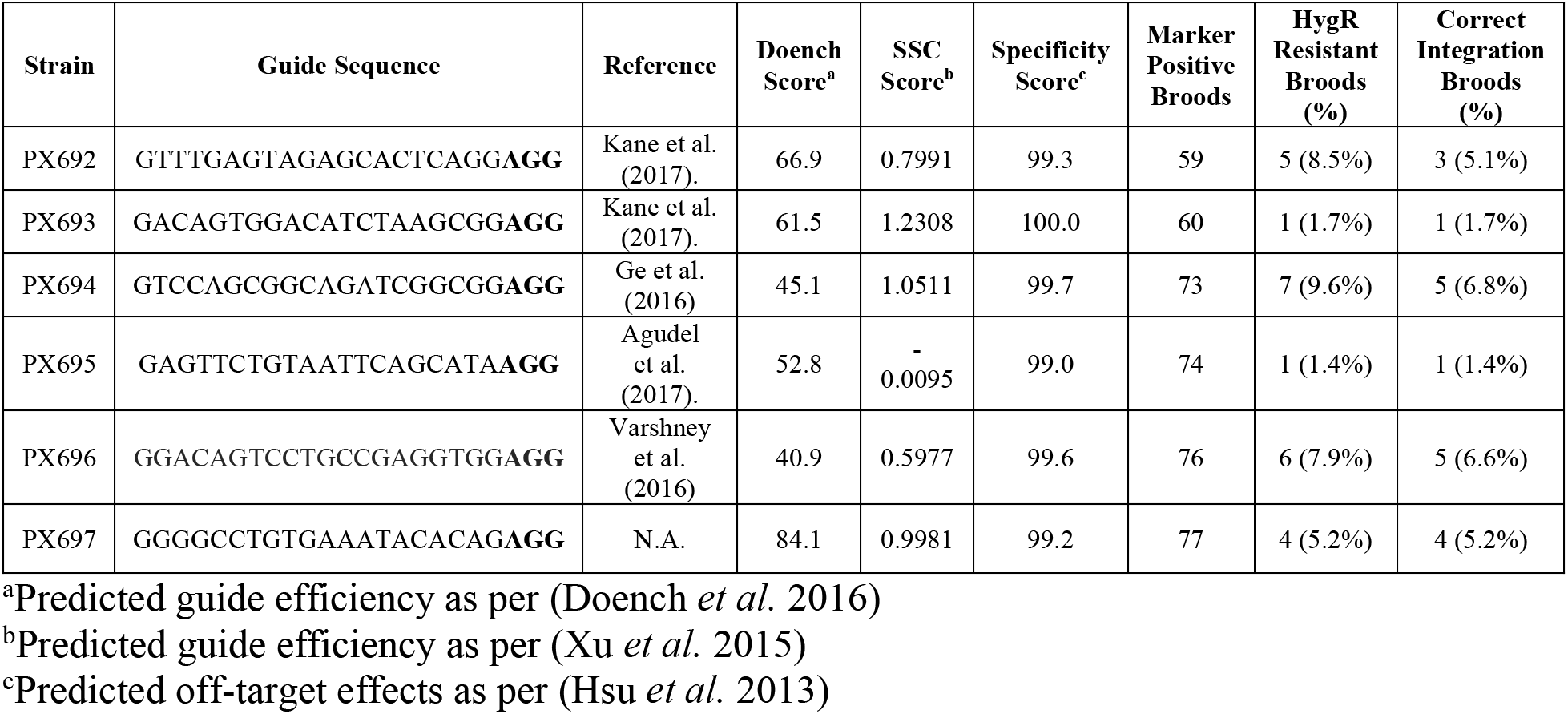
SLP guide efficiency for insertion of *rpl-28p::mKate2*.

Repair template plasmids were assembled from overlapping fragments using the NEBuilder HiFi Kit (NEB) per manufacturer instructions. For the landing pads, the *Cbr-unc-119* rescue gene and a portion of the homology arms containing the guide site were removed from pCFJ151(Addgene #19330) (Frøkjær-Jensen *et al.* 2008) and replaced with a multiple cloning site to create pMS2. The SEC from pDD285 (Addgene #66826) (Dickinson *et al.* 2015) was then inserted into SacI digested pMS2 to create pMS4. The C-terminal portion of the hygromycin resistance gene and the *unc-54* 3’ UTR were then independently amplified from pCFJ1663 (Addgene #514840 from the lab of Erik Jorgensen) and inserted into the SbfI site of pMS4 to create the final landing pad plasmids (pMS70-75). The six synthetic guide sites were included in the primers used to amplify the hygromycin resistance fragment (Table 1, Table S2, and Table S3). A complete annotated sequence of pMS74 can be found in Figure S1.

To generate an *rpl-28p::mKate2::unc-54 3’UTR* reporter, *rpl-28p* amplified from pBCN39-R4R3 (Addgene #34914) (Semple *et al.* 2012) and the mKate2 coding sequence and *unc-54* 3’UTR amplified from pDD285 were inserted into SacI digested pMS2 to give pMS12. The reporter was then amplified from pMS12 and inserted into an intermediate construct containing: *rps-0p* and the N-terminal fragment of the hygromycin resistance gene from pCFJ1663 (amplified in two fragments to remove intron), a pUC57 backbone, a truncated 5’ genomic homology arm from pMS2 and artificial sequences; to give the final insertion vector pMS81. A complete annotated sequence of pMS81 can be found in Figure S2. A second split hygromycin insertion vector, pZCS52, was made by amplifying the homology arms and split hygromycin from pMS81 by PCR and adding the *sqt-1(e1350)* gene amplified from pDD285.

To generate an additional fluorescent co-marker, *eft-3p* and *tbb-2* 3’ UTR amplified from pDD162 and wrmScarlet amplified from pSEM89 (Bindels *et al.* 2016; Mouridi *et al.* 2017) were cloned into a pUC19 backbone to give pZCS16. The CRE expressing plasmid pZCS23 was made by PCR amplifying the backbone, *eft-3p* and *tbb-2* 3’ UTR from pZCS16 and adding *NLS::CRE* from synthetic gBlocks (IDT).

### Strain generation by CRISPR/Cas9

A mixture consisting of 50 ng/μl pMS8, 10 ng/μl of the appropriate landing pad plasmid and 2.5 ng/μl pCFJ421 (Addgene #34876) (Frøkjær-Jensen *et al.* 2012) was microinjected into the gonad of young adult N2 hermaphrodites. Screening and removal of the SEC were done following Dickinson *et al.* (2015). Presence of the insertion and removal of the SEC was confirmed by PCR and Sanger sequencing (Table S3). Confirmed transgenics were backcrossed once to N2 to create the final strains PX692-PX697 (Table 1, Table S1).

### Quantification of synthetic guide RNA efficiency

For each landing pad strain (PX692-PX697), a mixture consisting of 50 ng/μl of all-in-one plasmid targeting the corresponding synthetic guide site and 10 ng/μl pMS81 was microinjected into the gonad of young adult hermaphrodites. Following injection, all worms were maintained at 25°C for the duration of the experiment. Injections were performed until approximately 60 broods per strain had at least one F1 progeny expressing the fluorescent donor homology, thereby marking the brood as successfully injected. Broods were screened for fluorescence at approximately 48h post-injection (hpi), and all fluorescent individuals were counted regardless of developmental stage (Figure S3). Hygromycin B was then top spread to plates at a final concentration of 250 μg/ml and plates were then screened starting five days later for resistant progeny. Individuals from surviving broods were PCR screened to confirm correct integration.

### Removal of the hygromycin selectable marker with CRE

A confirmed homozygous integrant line for *rpl-28p::mKate2::unc-54 3’UTR* was injected with 10ng/μl of CRE expression plasmid pZCS23 and 10ng/μl pCFJ421 co-marker. 30 co-marker positive F1 individuals were screened by PCR for the removal of the hygromycin gene. F2 progeny from 3 of the most promising candidates were then rescreened to confirm homozygous removal.

### In-situ assembly for integrated transgenes

Two or six PCR fragments with 30bp overlaps, covering the *sqt-1(e1350)* gene, were amplified from pDD285 using Q5 polymerase (NEB) per manufacturer instructions. Homology arms with 30bp overlaps to the *sqt-1(e1350)* gene were similarly amplified from pMS81. These homology arms were then complexed with the adjoining *sqt-1(e1350)* PCR fragment through a second round of PCR.

For *in-situ* assembly and integration, a mixture consisting of 50ng/μl pMS79, 5ng/μl pZCS16, and 40fmol/μl of each of the appropriate gel purified PCR products was microinjected into the gonad of young adult PX696 worms. As a control, 10 ng/μl pZCS52 was substituted for the PCR products. Following injection, all worms were maintained at 25°C for the duration of the experiment. After 24 hours, injected adults were moved to new plates to facilitate counting. F1 individuals were screened for red fluorescence (Figure S4) and the roller phenotype at 3-4 days post-injection. Hygromycin B was then added to plates at a final concentration of 250 μg/ml. Each day for five days post-exposure, plates were scored for hygromycin resistance. Individuals resistant to hygromycin and with the roller phenotype were singled without hygromycin and screened for Mendelian inheritance of the roller phenotype to indicate an integration event. Lines with promising candidates for single copy-integration were singled until they produced homozygous *rol* progeny, which were then screened for the presence of the wrmScarlet co-marker, genotyped by PCR across the insert and Sanger sequenced for correct transgene assembly and integration (Table S3).

### Accessibility of reagents and protocols

pMS4, pMS74, PMS79, pMS81, and pZCS16 are available from Addgene. Strain PX696 is available from the *Caenorhabditis* Genetics Center. Other strains and plasmids are available upon request. Full protocols and all plasmid sequences are available on the lab website (github.com/phillips-lab/SLP). All data files associated with this manuscript are available on FigShare.

## Results

### Generation of synthetic split landing pad sites

Current methods of transgene insertion in *C. elegans* result in a large number of false positives requiring additional phenotypic or PCR screening or anti-array selection. We sought a faster and simpler method by using CRISPR/Cas9 genome engineering to custom-designed synthetic split landing pads (SLPs). A split uracil selection system has been developed for yeast, whereby a functional *URA3* gene is reconstituted by an integration event. Therefore only transgenic cells survive in the absence of uracil (Levy et al. 2015). We reasoned a similar system could be adapted to *C. elegans*. Hygromycin resistance was chosen as the selectable event as it works across *C. elegans* strains, provides more substantial selection than other antibiotic resistance genes, and does not rely on mutant backgrounds such as with *unc-119* rescue (Radman *et al.* 2013). An artificial guide site plus the 3’ hygromycin coding sequence and transcription terminator were integrated into the target genome site as part of the SLP, whereas the promoter and 5’ portion of the hygromycin coding sequence were included in the repair plasmid (Figure 1A). A central 500bp region was included in both fragments, allowing for homology-directed repair. Since a complete resistance gene is not present in either the insertion strain or extrachromosomal arrays containing the repair plasmid, only individuals with proper homology-directed repair have a functional hygromycin resistance gene and survive hygromycin exposure (Figure 1B-D).

**Figure 1.**
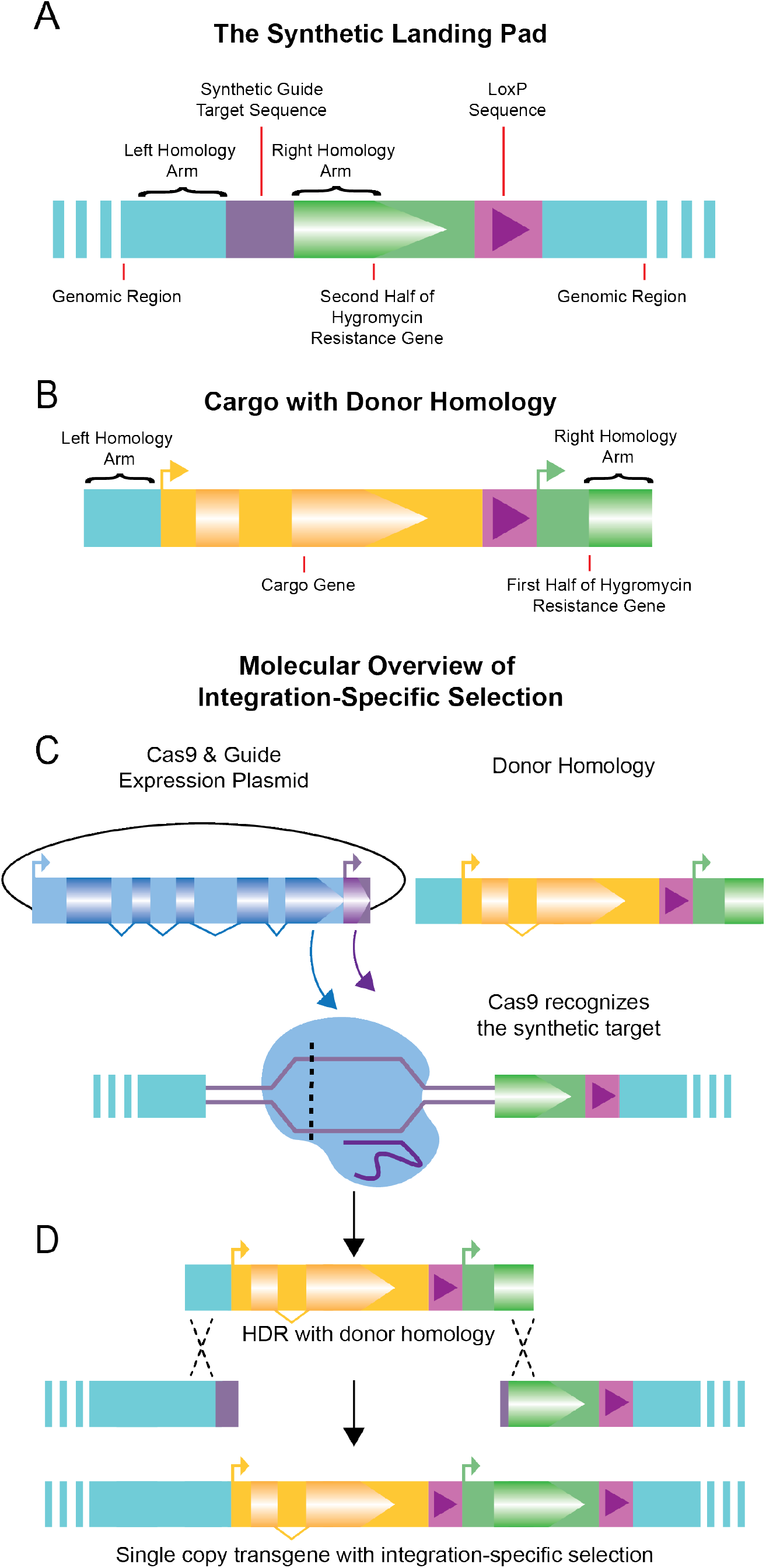
Overview of Integration Specific Selection. **A)** The Synthetic Landing Pad (SLP) with synthetic guide RNA target sequence, the 5’ fragment of the hygromycin resistance gene (partial coding sequence and UTR), and a single LoxP sequence. **B)** The donor homology with cargo transgene to be inserted, a second LoxP sequence, and the 3’ fragment of the hygromycin resistance gene (promoter & partial coding sequence). **C)** Cas9 & guide expression plasmid is injected with donor homology. Cas9 targets and creates a double-strand break at the synthetic target location. **D)** Once the double-strand break is made, repair with the donor homology integrates the transgenic cargo, and restores the hygromycin gene, allowing selection to occur only upon integration.

The SLP was inserted at Chromosome II:8,420,157. Both this general region and this specific CRISPR site have been shown to be permissive for gene expression, including germline expression (Frøkjær-Jensen *et al.* 2008, 2012; Dickinson *et al.* 2015). The SLP was introduced using the SEC selection method, and removal of the SEC left a LoxP site downstream of the hygromycin resistance gene terminator (Figure 1A). By also including a LoxP site upstream of the promoter in the repair template, this allows for optional removal of the HygR gene in confirmed integrants by injection of a CRE expressing plasmid (Figure S5).

### Efficiency of transgene integration

Given that the SLP is entirely artificial, the guide site can be of the experimenter’s choosing. As previous work has shown that the choice of guide site can influence integration efficiency (Farboud and Meyer 2015), we made six different SLP strains, each differing only in their guide site (Table 1). Five of these were sites previously shown to be highly efficient in other model organisms, while the sixth was designed to maximize the predicted guide efficiency. All had very low predicted off-target effects. We then attempted to integrate into each of these a universal *rpl-28p::mKate2* repair template (pMS81), which also served as a marker of injection success as it can also be incorporated into and expressed in extrachromosomal arrays. 1.4-9.6% of successfully injected broods (as determined by the presence of at least one mKate2 positive individual at 48 hpi) produced hygromycin resistant individuals. Overall, 79.2% of hygromycin resistant broods (19 of 24) also had perfect integrations events as determined by PCR (Table 1). Imperfect integrations are most often the result of HDR on one side of a double-strand break and incorrect integration on the opposite side, a known issue with transgenic integrations, although rearrangements within the transgene also occur, likely as a byproduct of the array assembly process (Stinchcomb *et al.* 1985).

Overall, we found three guide sites to have similar relatively high efficiencies, while one was slightly less efficient and the other two were much less efficient. The observed efficiencies were not always consistent with the predicted guide site efficiencies. For the top three guide sites 7.9-9.6% of co-marker positive broods contained integrants (not all of which were correct) which equated to approximately 300-450 co-marker positive progeny per integration event. In our hands, injection of thirty total individuals from any of the four best-performing strains was nearly always sufficient to obtain at least one correct line. This is on par with other transgene integration methods directed at this region (Frøkjær-Jensen *et al.* 2008, 2012; Dickinson *et al.* 2013, 2015) which have variable integration efficiencies depending on the transgene, but generally range from 5-30% with a few exceptions reaching higher frequency (Frøkjær-Jensen *et al.* 2012). As previously observed (Paix *et al.* 2014), broods with larger numbers of injection marker positive progeny (jackpots) were the most likely to yield integrants. However, this number was not predictive of perfect versus imperfect integration (Figure S3).

### In-situ donor assembly and integration

While plasmids offer the advantage of producing large quantities of the repair template, they require time and labor to produce. Standard cloning practices require a source of DNA, a ligation step, bacterial transformation, plasmid purification, and verification. As we sought to both simplify the process and reduce the overall time-to-integration, we attempted to bypass the cloning step and utilize the *C. elegans* native homology-directed repair to produce a transgene (Figure 2). While clone-free transgenesis has been previously demonstrated in *C. elegans* (Paix *et al.* 2016; Philip *et al.* 2019) we wanted to see if this process could synergize with the split-selection system to further improve the process as the previous work did not provide direct selection on the integration event. To test this approach, we utilized the *sqt-1(e1350*) mutation as it gives a dominant roller phenotype allowing us to assay for *in-situ* assembly. Transgene integration was most efficient for the plasmid vector, with 20% of co-marker positive broods containing an integrant. However, confirmed *in-situ* assembled and integrated *sqt-1* genes were also obtained using both two and six PCR fragments (Table 2, PCR Confirmed Integrations). Two parts correctly assembled and integrated more often than six parts. In some cases, hygromycin resistant individuals were observed without the *sqt-1* roller phenotype (Table 2, Hygromycin Resistant Broods). We believe these represent incorrect integration events, where at least the 5’ hygromycin resistance coding fragment was integrated into the genome. As these cannot be correct integration and assembly events, we did not pursue or characterize them.

**Figure 2.**
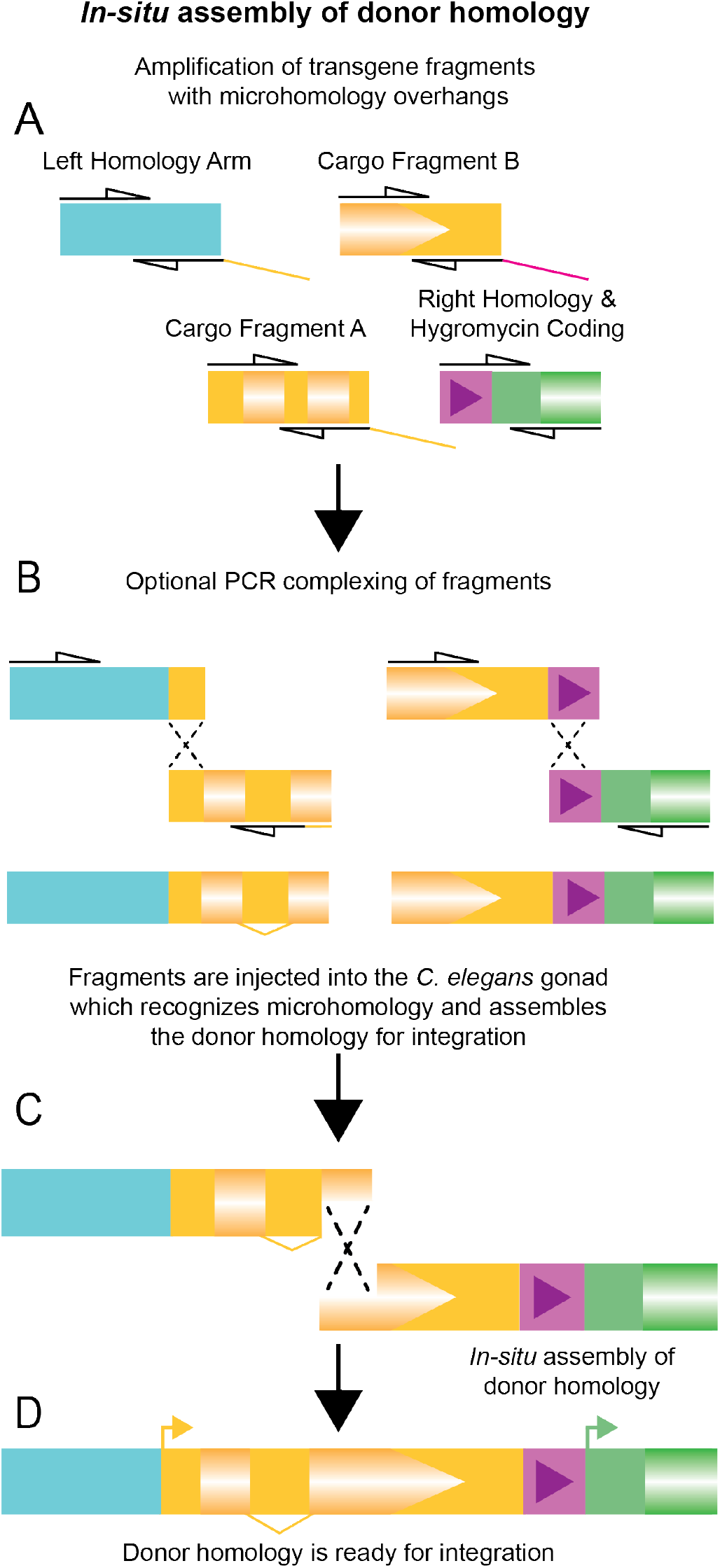
Overview of *in-situ* assembly. **A)** Amplification of homology arms and cargo fragments by PCR with overlaps of ~30bp **B)** Optional complexing by a second round of PCR reduces the number of fragments and increases the frequency of correct integration. **C)** Upon microinjection, PCR products are recombined by the worm using microhomology to make **D)** the complete donor homology ready for integration.

**Table 2:** In-situ assembly & integration efficiency.

|  |  | **Hygromycin Resistant Broods** |  |  | ***In-situ Assembly*** |  |  |
| --- | --- | --- | --- | --- | --- | --- | --- |
|  | **Marker Positive Broods** | **All Roller** | **Mixed** | **All wt** | **Homozygous Roller Isolated** | **PCR Confirmed Integrations** | **Error Free Integrations** |
| Plasmid | 15 | 3 | 0 | 0 | 3 | 3 | 3 (20.0%) |
| 2pc PCR | 41 | 6 | 3 | 0 | 8 | 6 | 3 (7.3%) |
| 6pc PCR | 51 | 0 | 8 | 5 | 1 | 1 | 0 (0.0%) |

For two-part assemblies, most hygromycin resistance events were accompanied by *sqt-1* assembly and integration, as indicated by the ability to isolate homozygous roller populations. However, not all of these insertions matched the expected sequence. Two of the eight insertions could not be amplified by PCR, suggesting larger scale errors, while three had point mutations identified during Sanger sequencing, and three had no detectable errors. In no case did we detect multiple copy insertions. In the six-part experiment, all resistant plates had non-roller (incorrect) integration events, with a few having roller individuals as well (Table 2, Hygromycin Resistant Broods). In most cases, a homozygous roller line could not be isolated, suggesting these individuals were the result of correctly assembled genes in arrays paired with incorrect integrations. In one case, a homozygous roller line could be isolated, indicating multiple integration events had taken place in that brood. In this case, the roller causing integration had a correctly assembled *sqt-1(e1350)* gene but also contained a second copy of one of the homology arms which was identified by Sanger sequencing.

The inclusion of a fluorescent co-marker allowed for monitoring of array loss in this experiment. Since there is no selection on the transgene containing plasmid, any arrays that form should be rapidly lost. As expected, prior to the addition of hygromycin, array positive individuals could be seen. However, none of the homozygous individuals isolated for sequencing (approximately 3-5 generations after injection) contained arrays, demonstrating arrays are indeed quickly lost in this system. Even so, it remains best practice to confirm array loss through either use of an array co-marker or a vector specific PCR performed in conjunction with the genotyping PCR.

## Discussion

We have created a fast and efficient strategy for integrating transgenes into the *C. elegans* genome that bypasses some pitfalls and laborious steps present in other methods. Combining split selection with self-assembly of repair templates takes what before would require at best two to three weeks down to as little as a week, while simultaneously reducing the required expertise and overall hands-on time (Figure 3). The core technology relies on integration-specific selection, made possible by SLPs at a safe harbor insertion site. These SLPs can be inserted into any *C. elegans* strain using a single set of reagents since the protocol presented does not rely on a particular genetic background. The landing pad presented is universal, and the experimenter can choose any type of cargo to be integrated. However, the SLP design could be modified with additional elements to facilitate specific types of insertions, such as reporter constructs or allelic variants. While we see no obvious phenotypic effects from the constitutive expression of hygromycin resistance protein, the SLP includes LoxP sites, allowing for a second injection of a CRE expression plasmid to remove the hygromycin gene. While removal by CRE injection is relatively efficient (Figure S5), if routine removal of hygromycin is desired, incorporation of inducible CRE and a marker gene into either the landing pad or insertion vector would allow for self-excision using a protocol similar to SEC (Dickinson *et al.* 2015). Further, additional SLPs using either the same or different guide and selective gene could be inserted into other sites in the *C. elegans* genome known to be permissive for transgene expression. This would facilitate the construction of more complex, multigene transgenic nematodes. The antibiotics neomycin (Giordano-Santini *et al.* 2010), puromycin (Semple *et al.* 2010), blasticidin (Kim *et al.* 2014) and nourseothricin (Obinata *et al.* 2018) have all been used for selection in *C. elegans,* and the coding sequences of the corresponding resistance genes could be split. The SLP, conceptually adopted from yeast, does not need to be restricted to *C. elegans.* While the formation of heritable arrays is unique to *Caenorhabditis* nematodes among model systems, and thus does not complicate single-copy integration transgenesis in other models, the concept of custom-designed SLPs could provide direct readouts for integrations events, with specific, custom-built guide target sequences. For example, a split fluorescent coding sequencing could suffice to screen injected embryos for proper, site-specific integration in model vertebrates and be coupled with a non-native, experimentally chosen guide RNA to reduce off-target effects while increasing on-target cutting and HDR.

**Figure 3.**
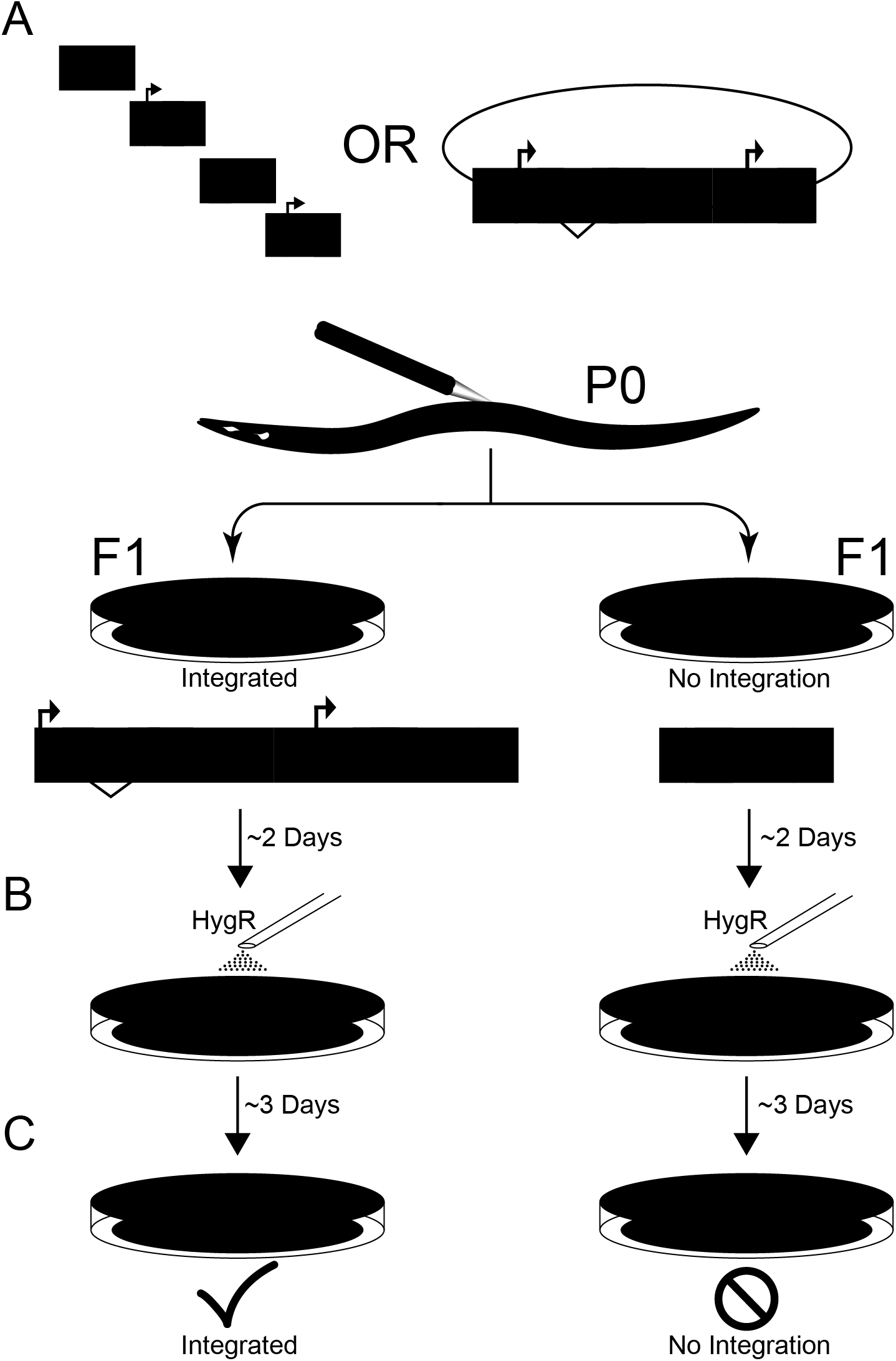
Experimental Overview. **A)** Injections of PCR fragment or cloned donor homology with Cas9 and guide expression plasmid. With PCR products, *in-situ* assembly forms the cargo transgene. **B)** Two days after injection, worms are exposed to hygromycin B. Since the array does not provide selection **C)** only integrated worms survive the exposure, providing the integration-specific selection.

The ability of *C. elegans* to self-assemble exogenous DNA fragments based on microhomology represents a possible alternative to plasmid cloning for insert assembly. Individuals with an assembled and integrated transgene were seen using both two or six PCR products. However, six pieces resulted in fewer correct integration and more incorrect integrations (Table 2, Hygromycin Resistant Broods), suggesting that, as expected, proper assembly occurs more often with fewer PCR products. As such, it is likely desirable to complex PCR products by overlap PCR before integration where feasible. While the use of PCR products, rather than plasmids, represents a more rapid protocol with fewer technical steps, it comes with the trade-off of a lower frequency of correct integrations requiring injection and screening of a larger number of worms. Thus, while use of PCR products results in a shorter time to integration confirmation, the total amount of hands on time is similar between the two protocols. Ultimately, at this time, choice of PCR products versus plasmid will largely come down to lab preference, although certain protocols, such as insertion of a library of similar constructs may favor the PCR approach.

Site-specific transgene insertions rely on homology-directed repair (HDR). However, non-homologous end joining is almost always the prevalent pathway in repairing a double-stranded break (Ward 2015; Xu *et al.* 2015). During guide efficiency testing, the top three guides all had similar insertion frequencies, suggesting we are approaching the upper limit of cutting efficiency and that further improvements will require improved rates of HDR. Furthermore, improved HDR should assist in the assembly of PCR products in worms and reduce the rate of false positives due to incorrect assembly. Low rates of HDR are not specific to *C. elegans* and impair HDR-based insertion in many model systems. As a result, multiple HDR enhancement strategies have been proposed in multiple model organisms (Beumer *et al.* 2008; Böttcher *et al.* 2014; Ward 2015). Adaptation and advancement of one or more of these methods will likely represent the next breakthrough in genome editing efficiency in *C. elegans*.

## Acknowledgments

This project was supported by grants from the National Institutes of Health (R01 AG56436, R35 GM131838) to PCP. We would like to thank Erin Petruccione for technical assistance and Stephen Banse for helpful discussions and comments on the manuscript.

---

**Figure S1.**
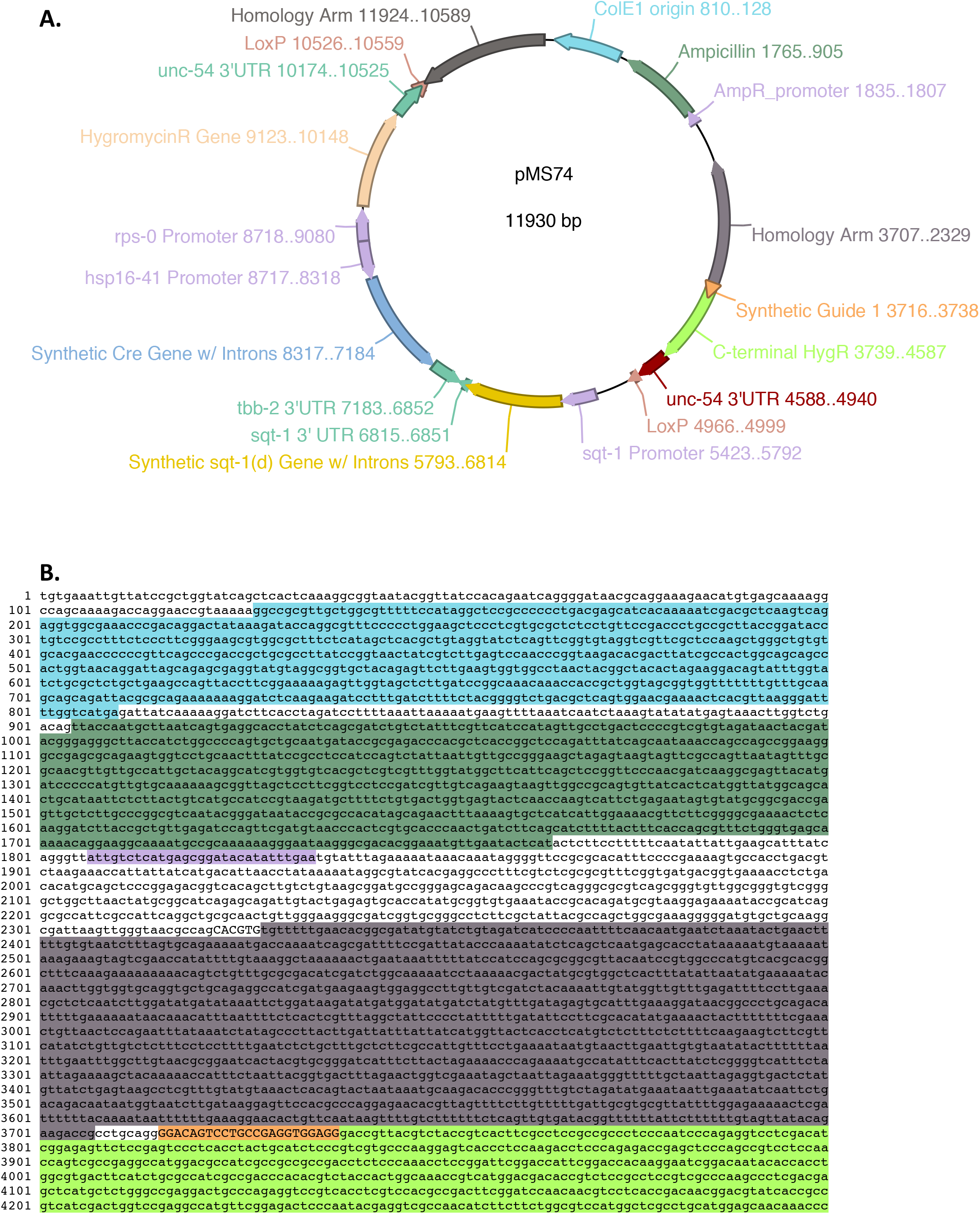
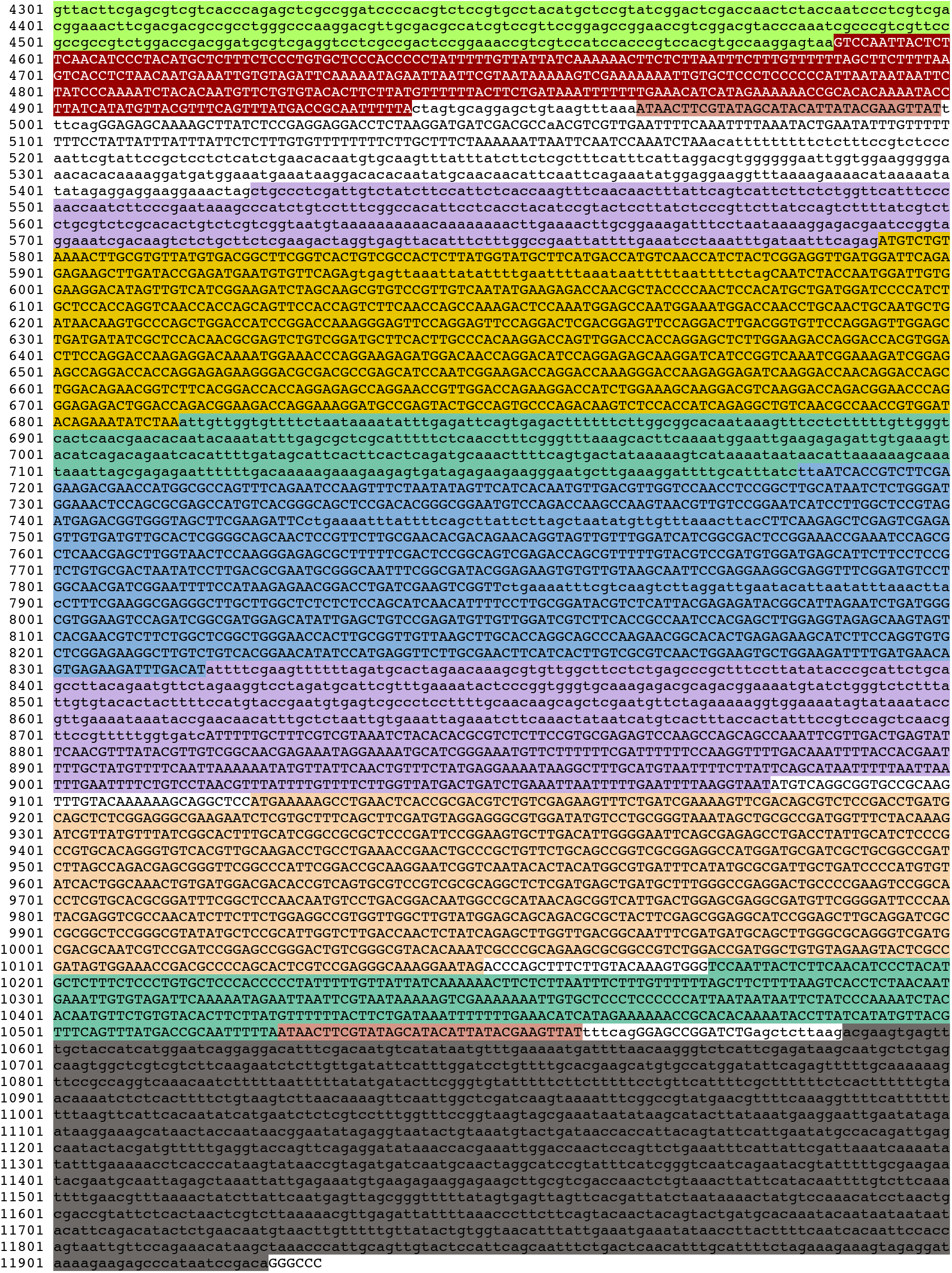
(A) Schematic of landing pad insertion vector pMS74 and (B) corresponding color-coded sequence

**Figure S2.**
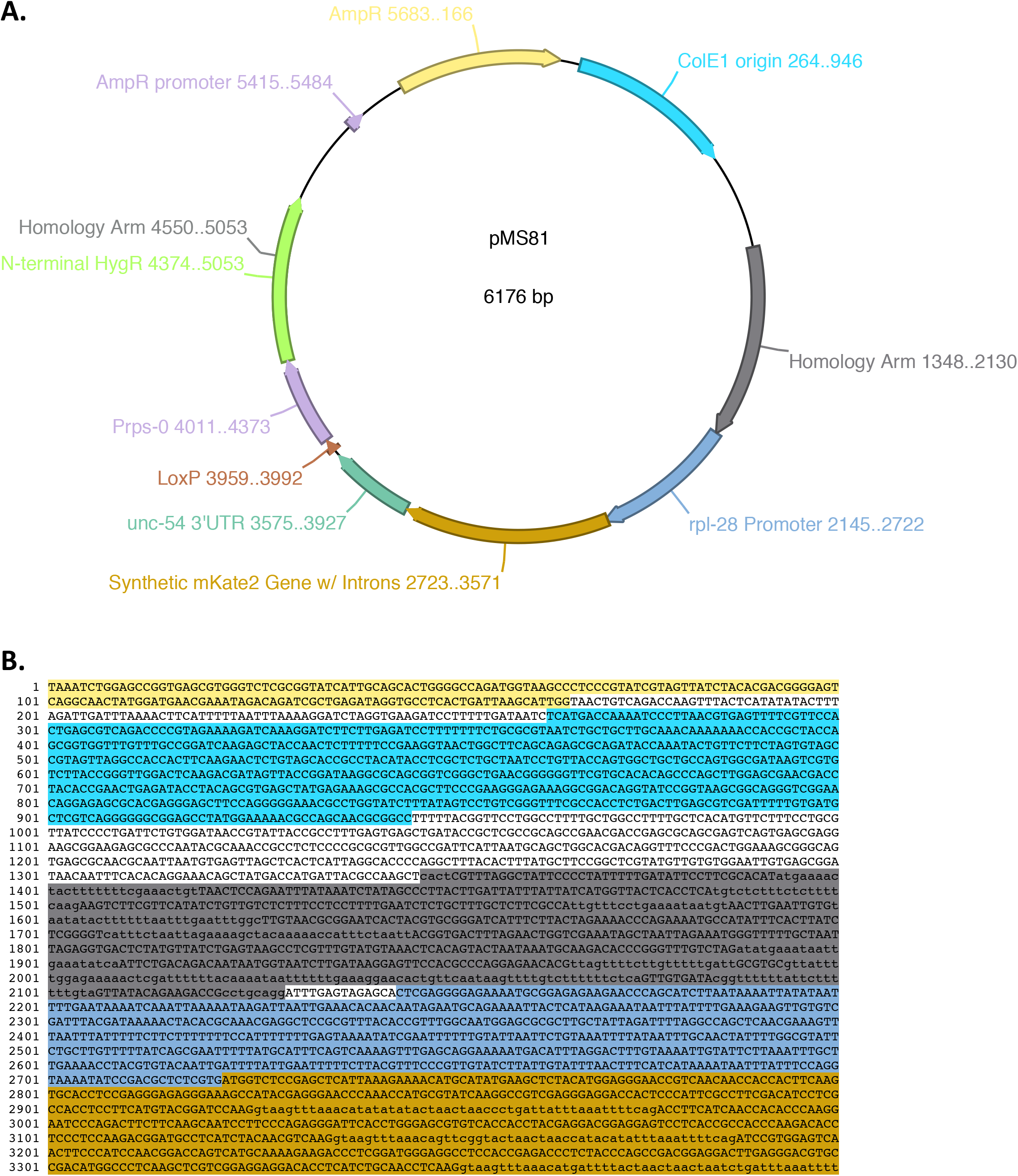
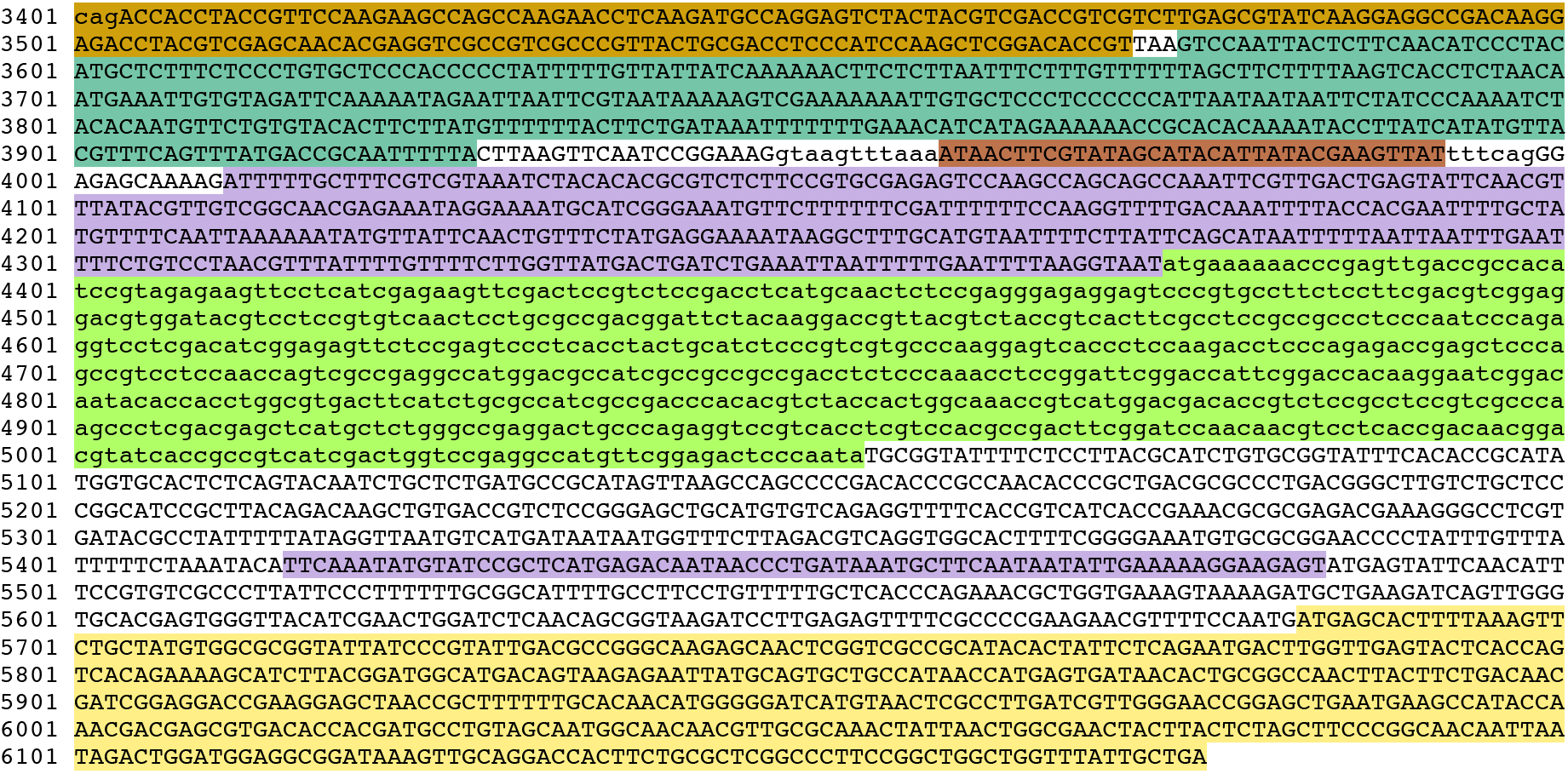
(A) Schematic of cargo insertion vector pMS81 and (B) corresponding color-coded sequence

**Figure S3.**
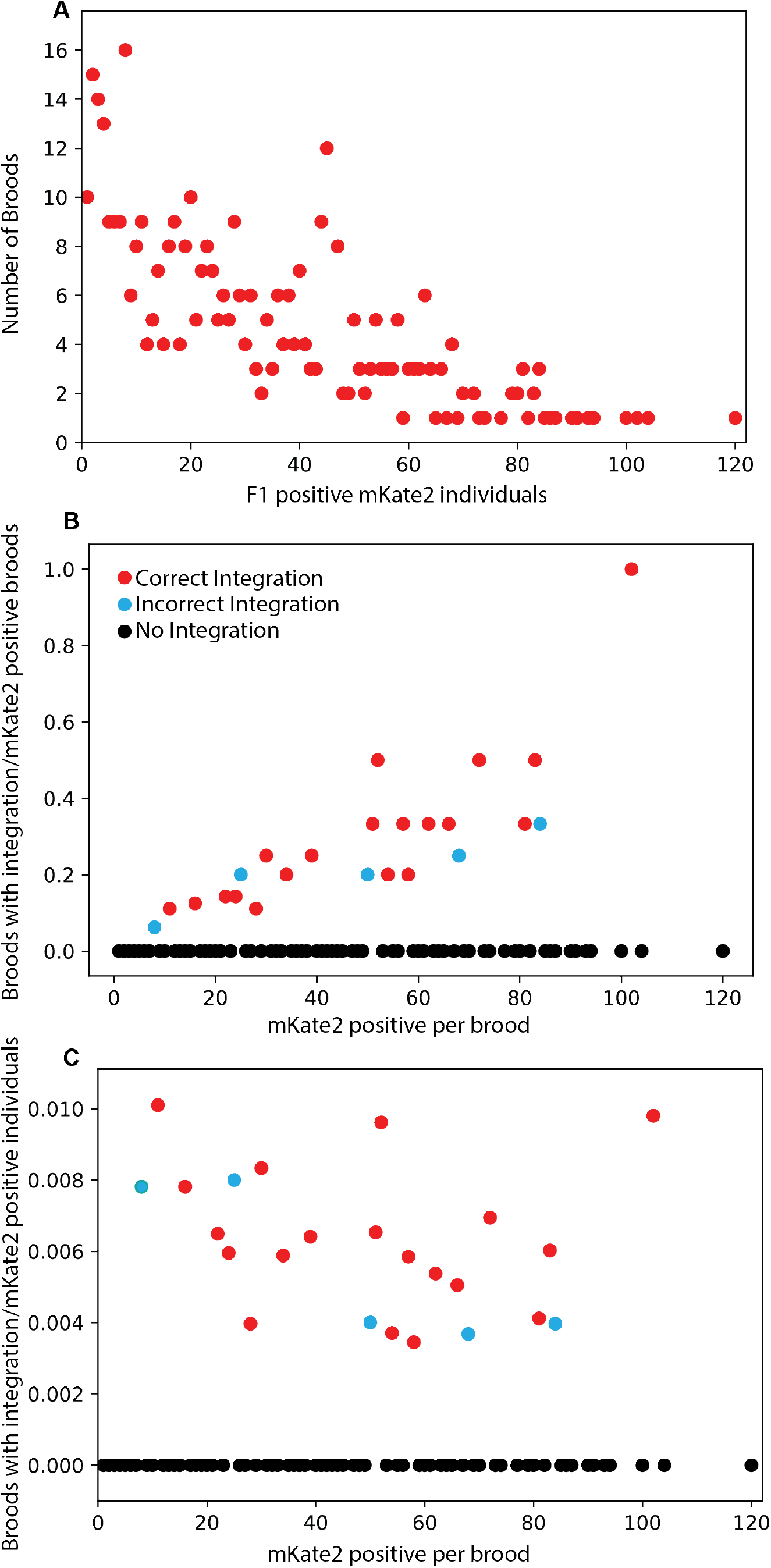
Comparison of integration broods. **A)** Distribution of injection marker *(mKate2’)* positive broods based on the number of *mKate2* positive F1 individuals per brood. Broods with a small number of positive individuals are most common; however, there are ‘jackpot’ injections showing large numbers of individuals. **B)** Broods with a large number of *mKate2* positive F1 individuals are more likely to have integration events, both correct and incorrect. However, when controlling for the total number of *mKate2* positive individuals, **C)** each brood with positive F1s is nearly equally likely to have integration events. Thus, while jackpot broods may have more transgenics, having more transgenics spread across several plates is similarly sufficient for identifying successful integration events.

**Figure S4.**
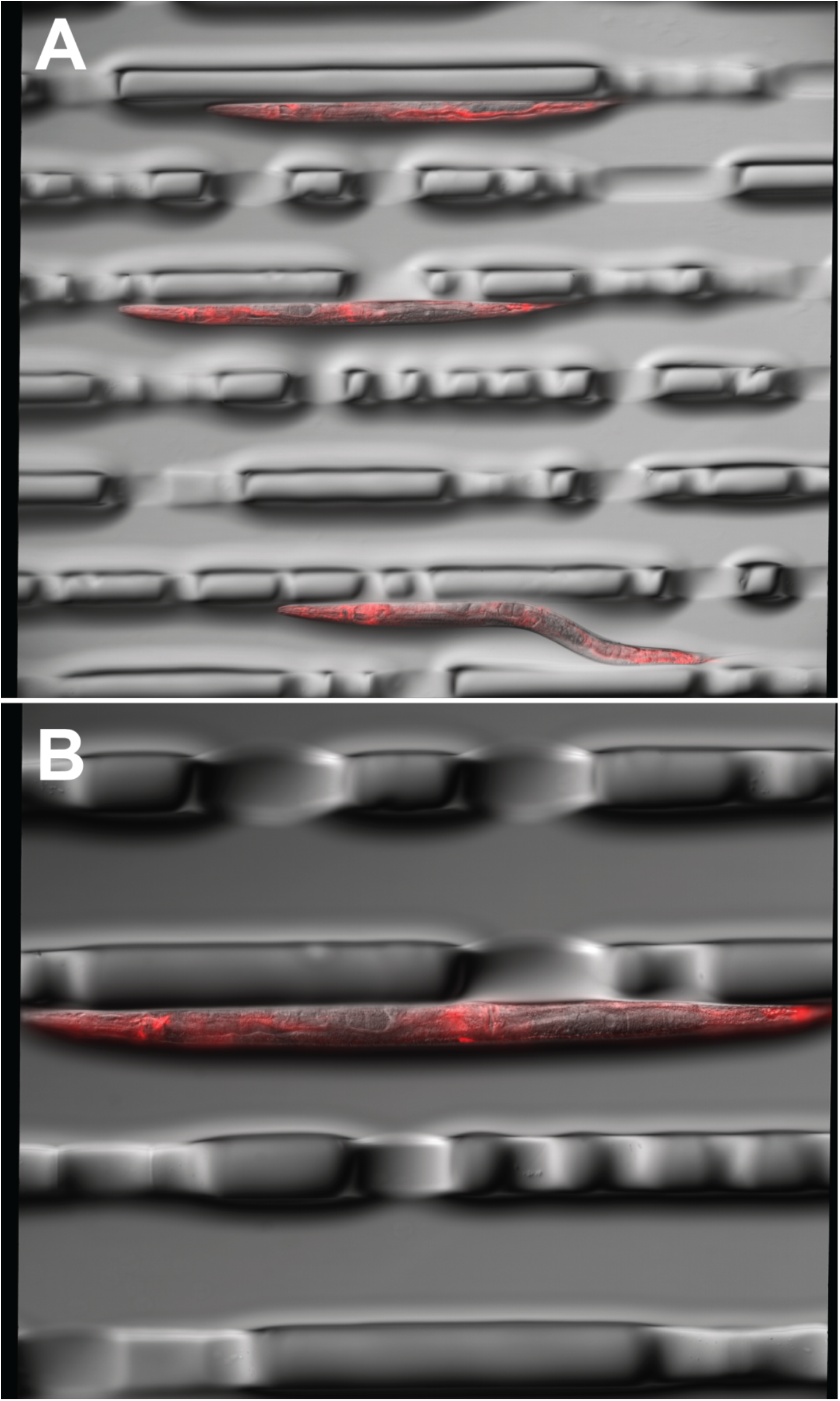
Representative images of *eft-3p::wrmScarlet::tbb-2 3’UTR* (pZCS16) array individuals. Synchronized young adults were injected with a mix containing 3 ng/μl pZCS16 91.25 ng/μl total DNA. A) Composite image of 30ms DIC and 40ms mCherry filter at 100x total magnification. B) Composite image 60ms DIC and mCherry filter at 200X total magnification.

**Figure S5.**
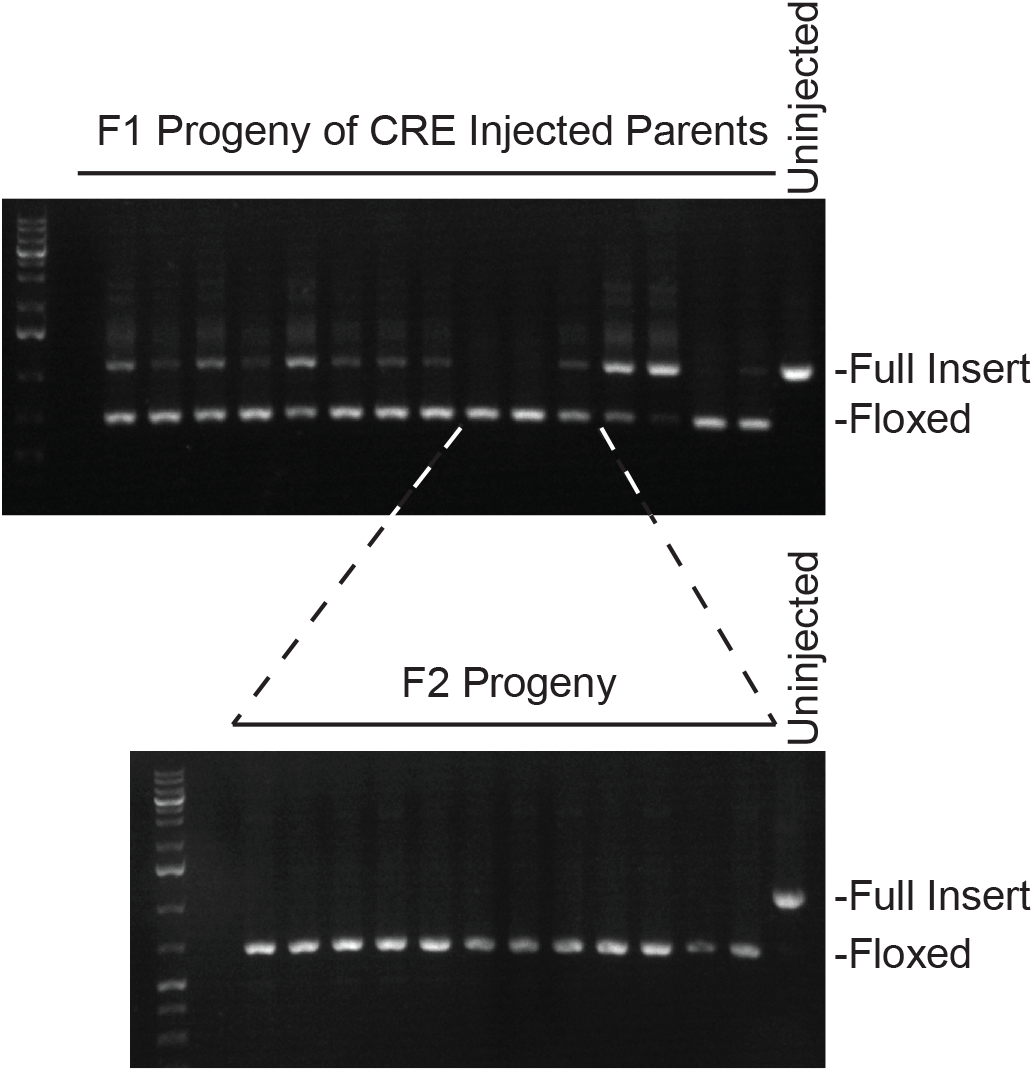
Injection of a CRE expressing plasmid results in the efficient removal of the hygromycin resistance marker gene. Co-marker expression marker positive F1 progeny (upper panel) were PCR screened for removal of the hygromycin resistance gene (“floxed” product). Four F2 progeny (lower panel) from three of the F1 broods with the weakest wild-type signal (“full insert” product), were then screened to confirm homozygous removal.

**Table S1.**
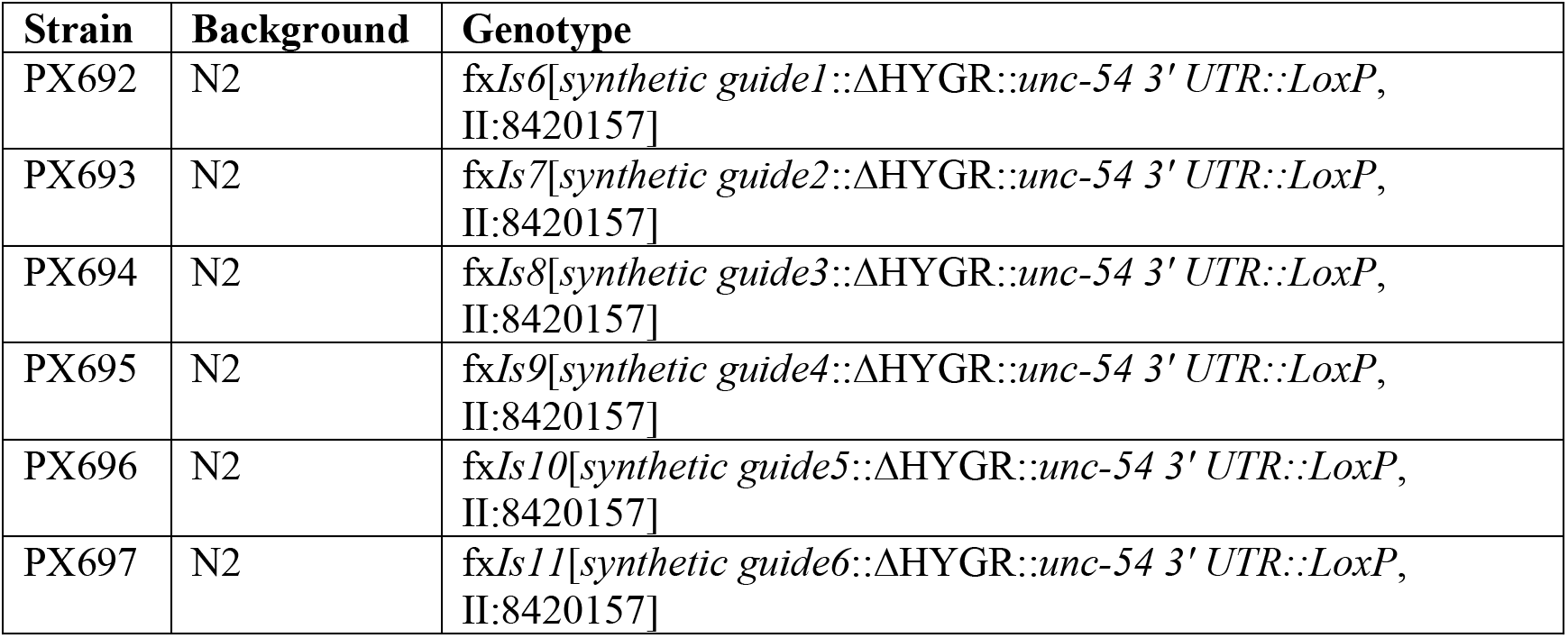
Strains generated for this study.

**Table S2.**
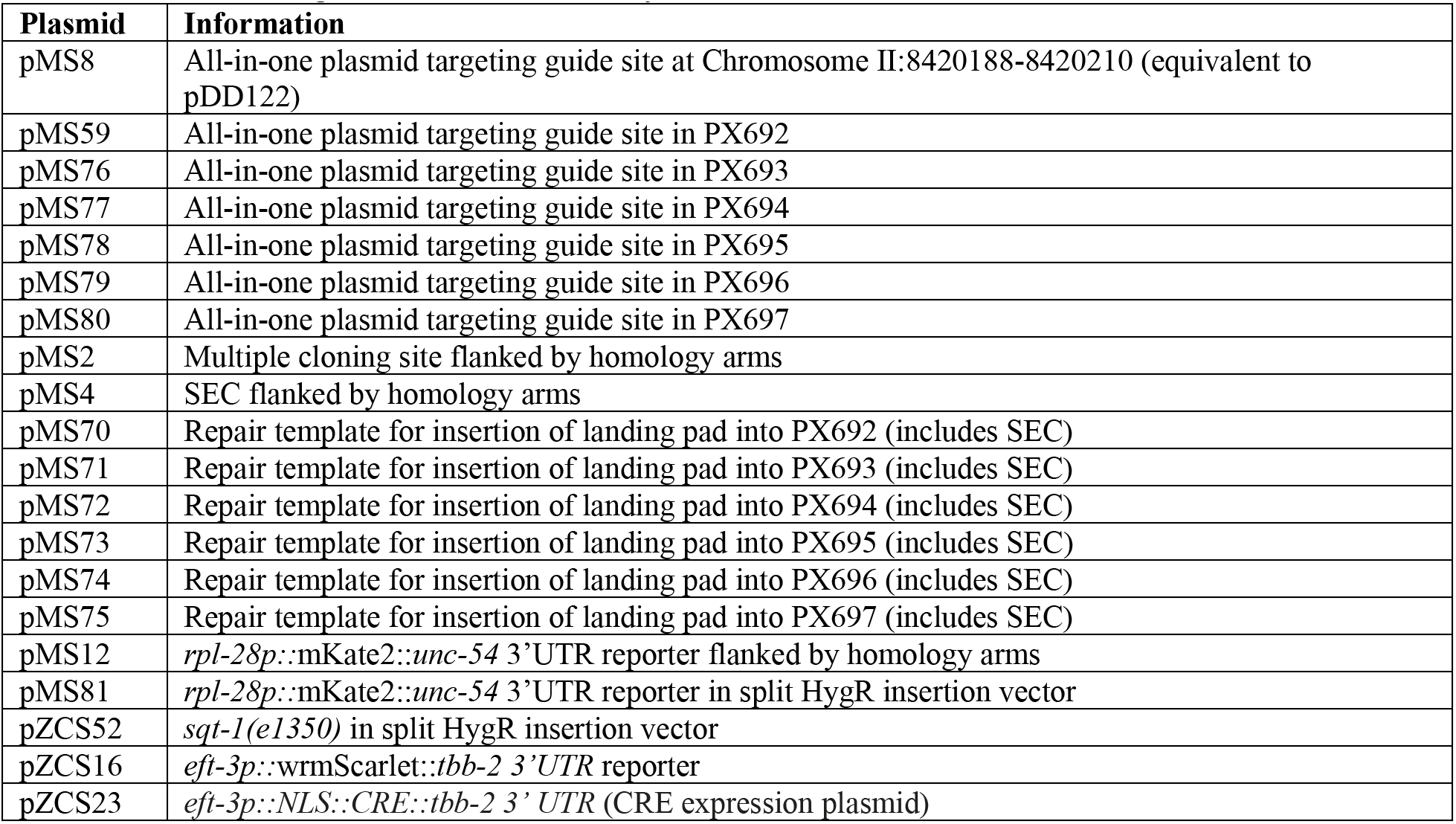
Plasmids generated for this study.

**Table S3.**
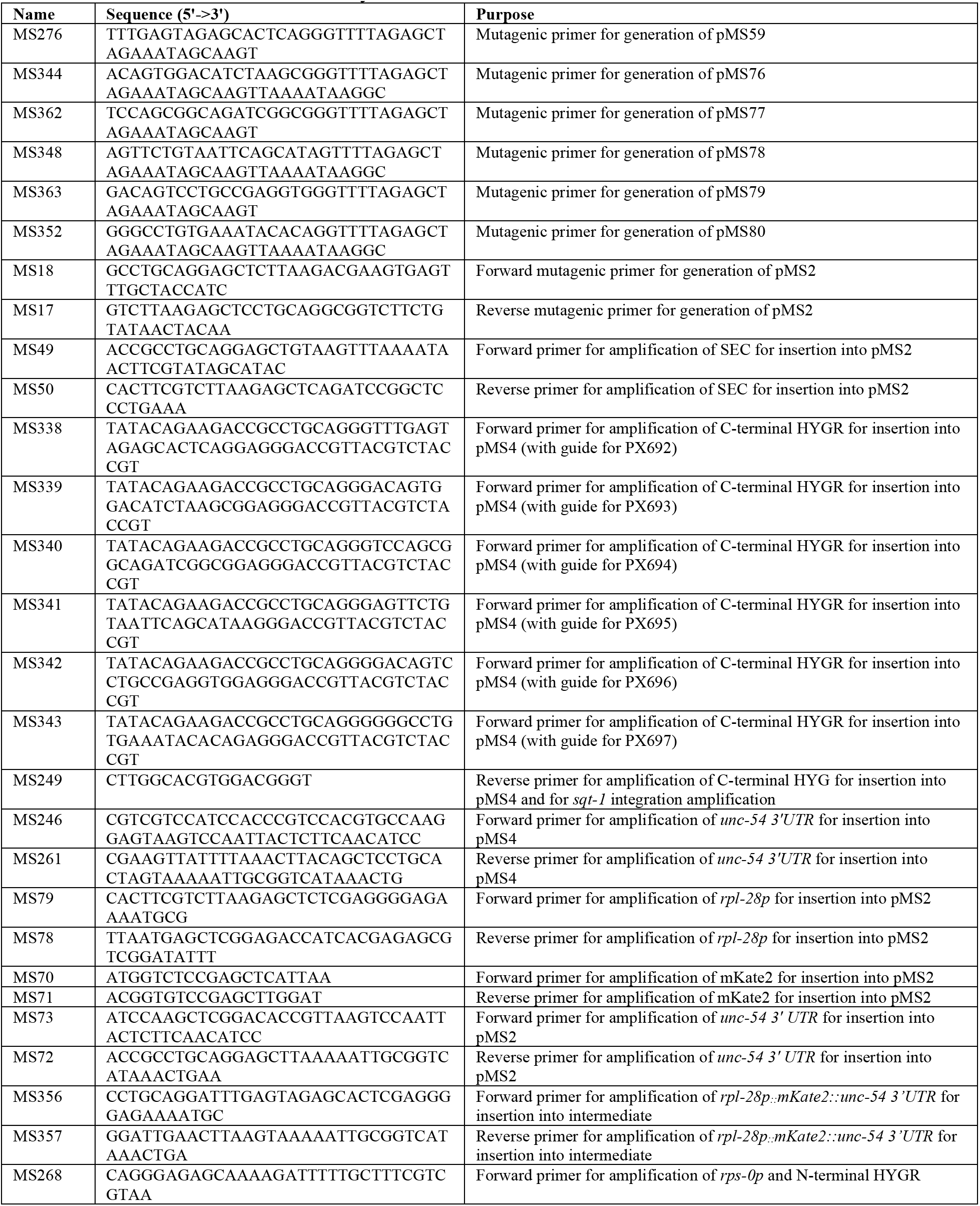
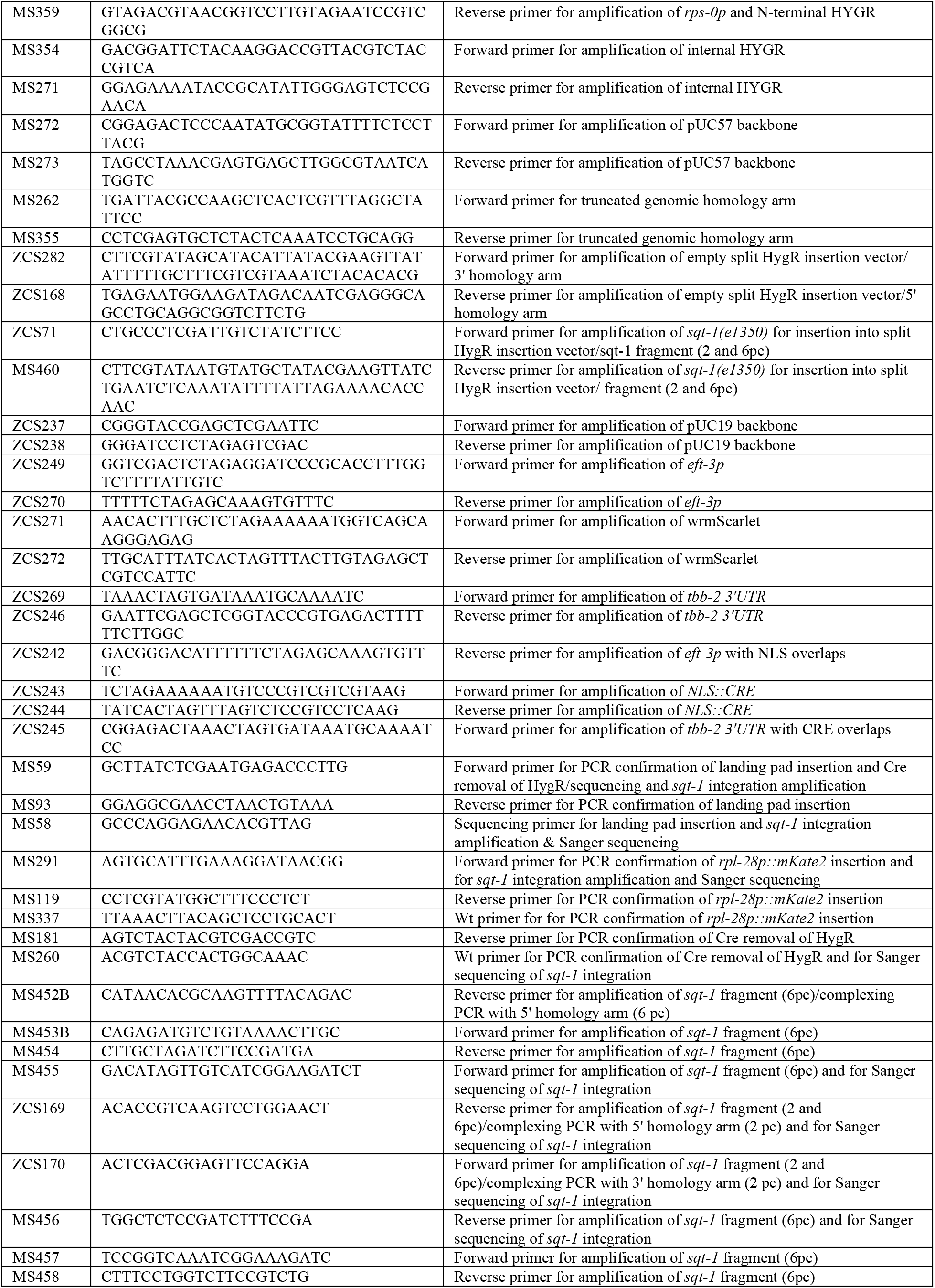
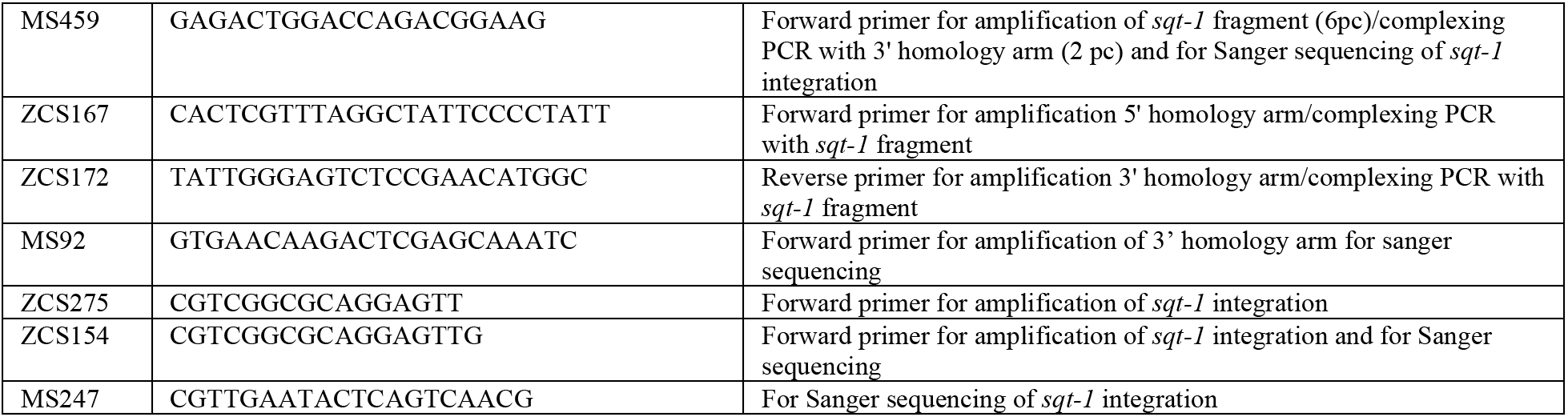
Primers used in this study.

